# Proteinarium: Multi-Sample Protein-Protein Interaction Analysis and Visualization Tool

**DOI:** 10.1101/589085

**Authors:** David Armanious, Jessica Schuster, George F. Tollefson, Anthony Agudelo, Andrew T. DeWan, Sorin Istrail, James Padbury, Alper Uzun

## Abstract

**Background:** Data analysis has become crucial in the post genomic era where the accumulation of genomic information is mounting exponentially. Analyzing protein-protein interactions in the context of the interactome is a powerful approach to understanding disease phenotypes.

**Results:** We describe Proteinarium, a multi-sample protein-protein interaction network analysis and visualization tool. Proteinarium can be used to analyze data for samples with dichotomous phenotypes, multiple samples from a single phenotype or a single sample. Then, by similarity clustering, the network-based relations of samples are identified and clusters of related samples are presented as a dendrogram. Each branch of the dendrogram is built based on network similarities of the samples. The protein-protein interaction networks can be analyzed and visualized on any branch of the dendrogram. Proteinarium’s input can be derived from transcriptome analysis, whole exome sequencing data or any high-throughput screening approach. Its strength lies in use of gene lists for each sample as a distinct input which are further analyzed through protein interaction analyses. Proteinarium output includes the gene lists of visualized networks and PPI interaction files where users can analyze the network(s) on other platforms such as Cytoscape. In addition, since the dendrogram is written in Newick tree format, users can visualize it in other software platforms like Dendroscope, ITOL.

**Conclusions:** Proteinarium, through the analysis and visualization of PPI networks, allows researchers to make important observations on high throughput data for a variety of research questions. Proteinarium identifies significant clusters of patients based on their shared network similarity for the disease of interest and the associated genes. Proteinarium is a command-line tool written in Java with no external dependencies and it is freely available at https://github.com/Armanious/Proteinarium.

## Background

Genome-wide association studies (GWAS) have become a popular approach to the investigation of complex diseases (1, 2) and have made possible discovery of insights not previously recognized (3–5). However, GWAS have often failed to demonstrate the “missing heritability” in many common diseases (6–10). A major factor contributing to the lack of success of genome-wide association studies in identifying ‘missing heritability’ is the fundamental nature of the genetic architecture interrogated by GWAS. The computational approaches underlying GWAS reflect the “common disease common variant hypothesis,” that complex disease architecture is due to additive genetic effects due to variants in individual genes. The genetics of complex diseases, however, suggests that is unlikely. The more likely architecture is that subgroups of patients share variants in genes in specific networks and pathways which are sufficient to give rise to a shared phenotype. It is also likely that variants in genes in different networks and pathways express similar phenotypes and define different subgroups of patients. All nodes in the pathways are unlikely to be equally represented. Because of purifying selection, highly expressed, pathogenic variants may be clustered in overlapping pathways, rate limiting steps or regions of high-centrality (11). Thus, resources are needed to identify these shared and individual networks and pathways in clusters of patients within diseases or phenotypes.

With the generation of high throughput screening methods, extensive protein– protein interaction (PPI) networks have been built (11, 12). PPI networks potentially harbor great power as they reflect the functional action of genes (13, 14). Large scale protein interaction maps show that genes involved in related phenotypes frequently interact physically at the level of proteins in model organisms, and as well as in humans (15–19). This is also true for the disease phenotypes. Proteins that are associated with similar disease phenotypes have a strong affinity to interact with each other (17, 20). In addition, they have a tendency to cluster in the same network zone (21). PPI networks have been used to identify candidate genes and subnetworks associated with complex diseases such as cancer and Alzheimer’s Disease (14, 22, 23).

We developed Proteinarium, a multi-sample protein-protein interaction tool to identify clusters of patients with shared networks to better understand the mechanism of complex disease and phenotypes. This tool was designed to increase the power in the analysis of experimental data by identifying disease associated biological networks that define clusters of patients, as well as the visualization of such networks with user specified parameters. Compared to other PPI network analysis tools, a distinguishing feature of Proteinarium is identification of clusters of samples (patients), their shared PPI networks and representation of samples within the clusters by their group assignment.

## Implementation

### Overview

Proteinarium is well suited for analyzing and visualizing the networks of complex diseases and their associated phenotypes to generate clusters of similar samples with their associated layered networks. Figure 1 presents an overview of the workflow for Proteinarium.

**Figure 1.**
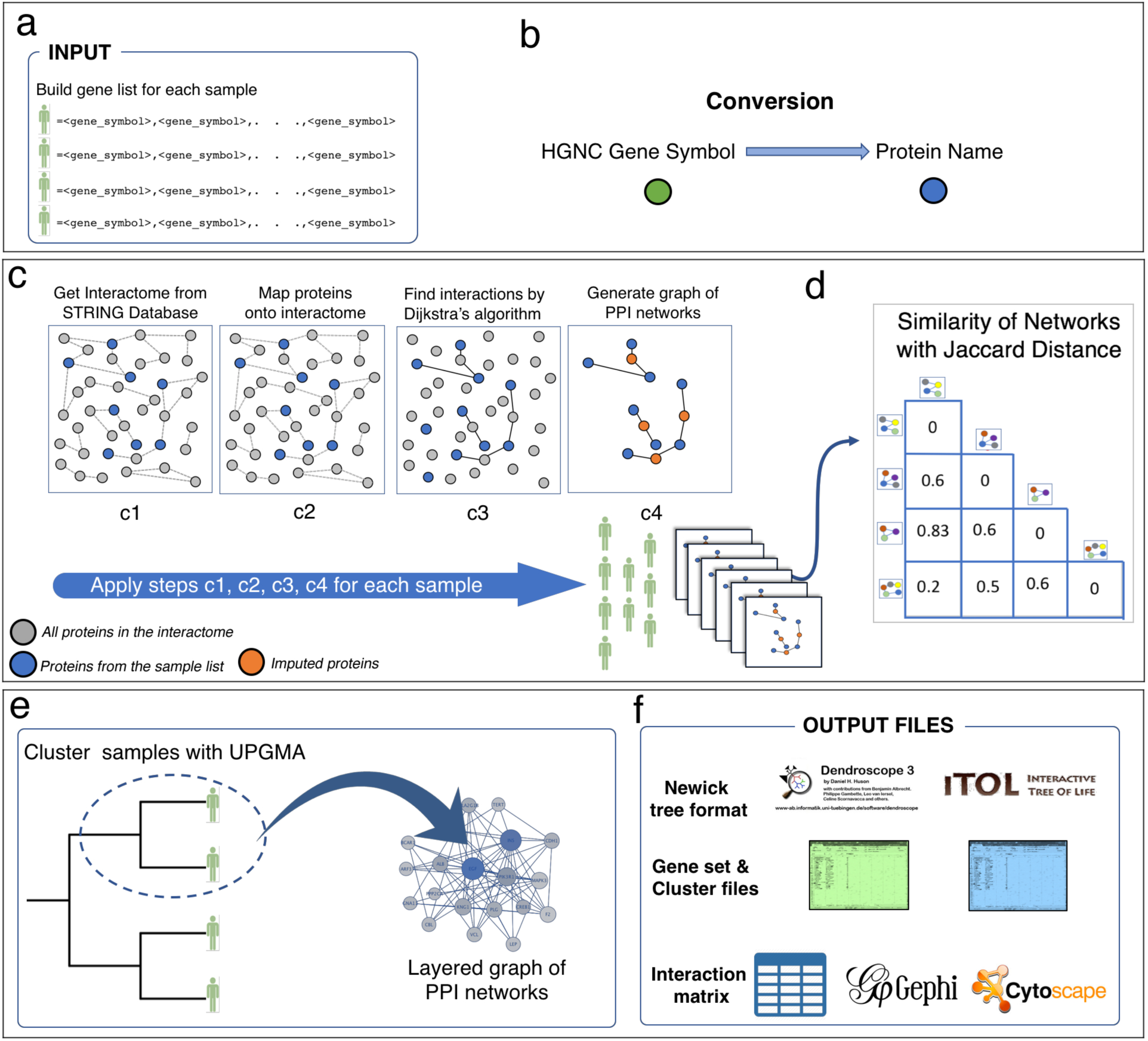
Proteinarium workflow. **(a)** Proteinarium accepts gene lists for each sample **(b)** All gene symbols are converted to their protein product names. **(c1)** Interactome Information is provided from STRING Database. **(c2)** Proteins for each sample are mapped onto interactome, red colored circles. **(c3)** Finding the interactions for each sample based on Dijkstra’s algorithm. **(c4)** Layered graph built for each sample based on their gene list. **(d)** Next the network similarity matrix between samples is built using Jaccard index. **(e)** Samples are clustered by UPGMA method based on their network similarities and any chosen branch of the dendrogram displays the layered graph of PPI networks which can be visualized on the fly and saved as an interaction file. **(f)**Proteinarium provides several output files for users to export the results to be used on other programs like ITOL, Gephi or Cytoscape.

### Data Source

Proteinarium uses the STRING database, version 11, for humans as its network data source. It includes known and predicted PPIs (7). Each PPI has an associated score between 0 and 1000 indicating the confidence of the interaction.

### Input Data

Proteinarium can be used to analyze data for samples with dichotomous phenotypes, multiple samples from a single phenotype or a single sample. For each sample, a list of genes is required as input (seed genes). If the samples are from a dichotomous phenotype, the genes for each sample must be in one of two discrete files, Group 1 Samples or Group 2 Samples. The seed genes may be derived from deep sequence data, transcriptome analysis, or from any analysis that generates candidate gene lists. Therefore, it is up to the investigator how to generate the seed genes as input for Proteinarium.

## Multi-sample Analysis: Constructing Networks

### Mapping onto the interactome

Proteinarium is initialized by mapping the seed genes onto STRING’s protein-protein interactome (19). For each seed gene, Proteinarium maps the HUGO Gene Nomenclature Committee (HGNC) Symbol to the associated proteins in STRING’s database (10). Proteinarium then finds all PPIs within the STRING database that correspond to the seed genes, forming a subset of the protein interactome for each sample.

### Building a graph with Dijkstra’s Algorithm

For each sample, Proteinarium builds an interaction graph from the seed proteins. We use Dijkstra’s shortest path algorithm to find short, high-confidence paths between all pairs of proteins where such a path exists (24). STRING provides a confidence score *S* for every edge connecting two proteins between 0 and 1000. The higher the number, the higher the confidence of the specific protein-protein interaction. We define the cost of an edge between two proteins to be 1000 − *S*. Thus, the highest confidence interactions should be the lowest cost edges. The algorithm has been modified to only consider paths whose lengths (number of vertices in the path) and total costs (sum of edge weights) are below the user specified values. For each sample *i*, the graph *G*_*i*_ = (*V*_*i*_, *E*_*i*_) is generated, where each vertex *v* ∈ *V*_*i*_ corresponds to a protein and each edge *e* ∈ *E*_*i*_ corresponds to a protein interaction. Only the seed proteins and the proteins that were required to minimally connect the seed proteins are included in the set of vertices *V*_*i*_.

### Building the similarity matrix and clustering samples with UPGMA

After generating the graphs for all samples, Proteinarium calculates the similarity between each pair of graphs using the Jaccard distance (25). The distance *d*_*i,j*_ between any two graphs *G*_*i*_ and *G*_*j*_ is calculated as:

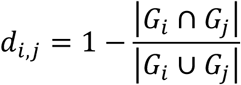

This similarity matrix is then used as the input to cluster the set of graphs. Clustering is performed hierarchically using Unweighted Pair Group Method with Arithmetic Mean (UPGMA) (12). Initially, all clusters consist of a single graph, corresponding to a leaf of a dendrogram. When there exists more than one cluster, we combine the two clusters that are closest to each other according the distance metric *d* and then update all cluster distances.

In the standard UPGMA algorithm, the weight *w*_*i*_ would correspond to the number of graphs in that cluster, so all the weights of all leaves in the tree would equal 1. Our modified algorithm scales either the Group 1 graph factor or the Group 2 graph factor to a constant *ρ* ≥ 1 such that the product of the size of the graph set and *ρ* is equal to the size of the other graph set.

## Visualization

### Visualizing the “Global Dendrogram”

Proteinarium outputs the results of the clustering algorithm as a dendrogram in both PDF visualization and in a text file in Newick tree format. The horizontal length of the line segments is proportional to the height of the cluster in the tree. The lower the height, the more similar the graphs within the cluster are. All heights are normalized between 0 and 1, with the leaves occupying height 0 and the ancestor occupying height 1. The lines are also colored according to the (weighted) percentage of Group 1 versus Group 2 gene sets comprising the cluster: if a group comprises more than 60% of the weight, then the edge is colored by that group; otherwise, the edge is colored black.

## Visualizing “Local”

### Clusters Constructing a layered graph

For a given cluster *C* comprised of *n* graphs *G*_1_ = (*V*_1_, *E*_1_),…, *G*_*n*_ = (*V*_*n*_, *E*_*n*_) Proteinarium can construct and output a summary layered graph *LG* = (*V*_*LG*_, *E*_*LG*_) consisting of each of the *n* graphs with the following vertices and edges:

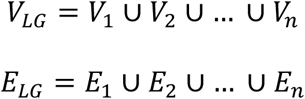

Additionally, the count annotation for each vertex *v* in *LG*:

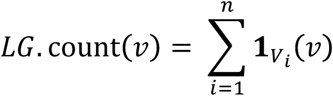

The count annotation is the number of sample networks in which the protein (v) of the layered graph is found. If a cluster *C* contains samples from both sample groups, there are 5 possible networks being created on the fly. These networks are as follows:

i. [Group 1] the network for Group1 samples only;
ii. [Group 2] the network for Group 2 samples only;
iii. [Group 1 + Group 2] the network for both Group1 and Group 2 samples;
iv. [Group 1 - Group 2] the network where Group 2 is subtracted from Group 1;
v. [Group 2 - Group1] the network where Group 1 is subtracted from Group 2.

Details on the constructions of graphs iv and v are in Supplementary Materials (S1a).

## Visualizing and annotating the layered graph

Given a layered graph *LG* corresponding to a cluster *C*, we lay the vertices according to a simple implementation of a force-directed layout algorithm. Additional details on the constructions of graphs iv and v are in Supplementary Materials (S1a). We color each vertex *v* according to the samples for which *v*’s corresponding gene exists:

1. Only samples from Group 1: yellow group1VertexColor
2. Only the samples from Group 2: blue group2VertexColor
3. Samples from both Group 1 and Group 2: green bothGroupsVertexColor or a 50/50 mixture of (1) and (2)
4. Neither samples from Group 1 nor Group 2 (Protein was inferred from the pairwise path finding algorithm): red [defaultVertexColor]

Additionally, the opacity of each vertex *v* is calculated linearly between a minimum opacity *m* and 255:

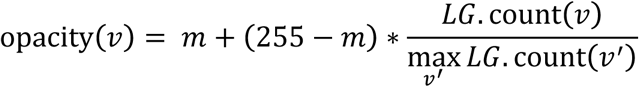

The same calculation is used for the opacity of the edges.

When a graph is to be visualized, the number of displayed vertices is reduced to keep the visualization tractable. We rank the seed genes of a given graph according to the number of pairwise paths that particular vertex appears in. We then retain the top *k* vertices such that the total number of unique vertices found within all pairwise paths of *k* selected vertices is less than or equal to the maximum number of vertices to be displayed. In doing so, all graphs will show only complete paths whose endpoints originate from seed genes.

### Fisher’s exact test p-value

The p-value for a cluster is given by a Fisher-Exact and indicates the probability of observing, among all *n* sample samples, a cluster of size *m* with proportions of Group 1 and Group 2 samples relative to the total number of Group 1 and Group 2 samples. In other words, it provides a sense of how disproportionate a particular group (either Group 1 or Group 2) is over-represented in a cluster; the lower the p-value, the more confidence we have that the over-representation is *not* due to random chance.

### Clustering coefficients

For each cluster, we calculate the global clustering coefficients for each of the four graphs: Group 1 graph, Group 2 graph, and the two graphs resulting from the weighted differences of each graph as described above. The clustering coefficient is a measure describing the tendency of vertices in a particular graph to cluster together. For a graph *G* = (*V, E*) with |*V*| = *n* vertices, the global clustering coefficient is given by

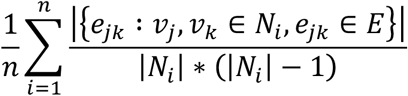

where *N*_*i*_ is the set of neighbors of vertex *v*_*i*_, i.e. *N*_*i*_ = {*v*_*i*_ ∶ *e*_*ij*_ ∈ *E*}.

### Program Availability

Proteinarium is a command-line tool written entirely in Java with no external dependencies. Java version 8 or above (Java 9, or Java 10) must be installed in order to run Proteinarium. It is freely available at https://github.com/Armanious/Proteinarium.

## Results

### Running Proteinarium on Simulated Data

We evaluated Proteinarium’s performance on simulated datasets and on the datasets for a use case. The STRING database was used to obtain “networks 1 and 2.” Both networks have total of 71 proteins and their clustering coefficients were respectively, 0.590 and 0.691. Members of the networks do not overlap. We simulated two groups each with 50 samples (Figure 2). For each sample in Group 1, 10 genes were randomly selected from PPI network 1 and for each sample in Group 2,10 genes were randomly selected from PPI network 2. We ran Proteinarium using these Group1 and Group2 seed gene files. To add noise to the simulated data, for each sample in Group 1 a percentage of their genes were randomly replaced with genes from PPI network 2 (20%, 30%, 40% and 50%). Similarly, for samples in Group 2, a percentage of genes were replaced at random with genes from PPI network 1. Representative dendrograms reflecting the output of Proteinarium for the simulations with 0%, 20%, 30%, 40% and 50% are shown in Figure 2 and the performance in the table below. Samples in Group 1 are colored yellow, and samples in Group 2 are colored blue. The five dendrograms show the clustering of the samples based on their network similarities. With no noise, as expected, samples from Group 1 perfectly cluster together and samples from Group 2 perfectly clustered together (power: 100%, p<.05) As more noise was added to the data sets, the power decreased from 100% for 20% noise, down to 15% for 40% noise. At 50% noise (i.e. the null hypothesis) the power to discriminated groups was less than 5%.

| Test | Replicate | Number of Simulations with P-value < 0.05 | Total Simulations | Fraction of Simulations with P-value < 0.05 | Number of Simulations with P-value ≥ 0.05 | Fraction of Simulations with P-value ≥ 0.05 |
| --- | --- | --- | --- | --- | --- | --- |
| 1 | 0.2 | 500 | 500 | 1 | 0 | 0 |
| 1 | 0.3 | 477 | 500 | 0.954 | 23 | 0.046 |
| 1 | 0.4 | 78 | 500 | 0.156 | 422 | 0.844 |
| 1 | 0.5 | 23 | 500 | 0.046 | 477 | 0.954 |

**Figure 2.**
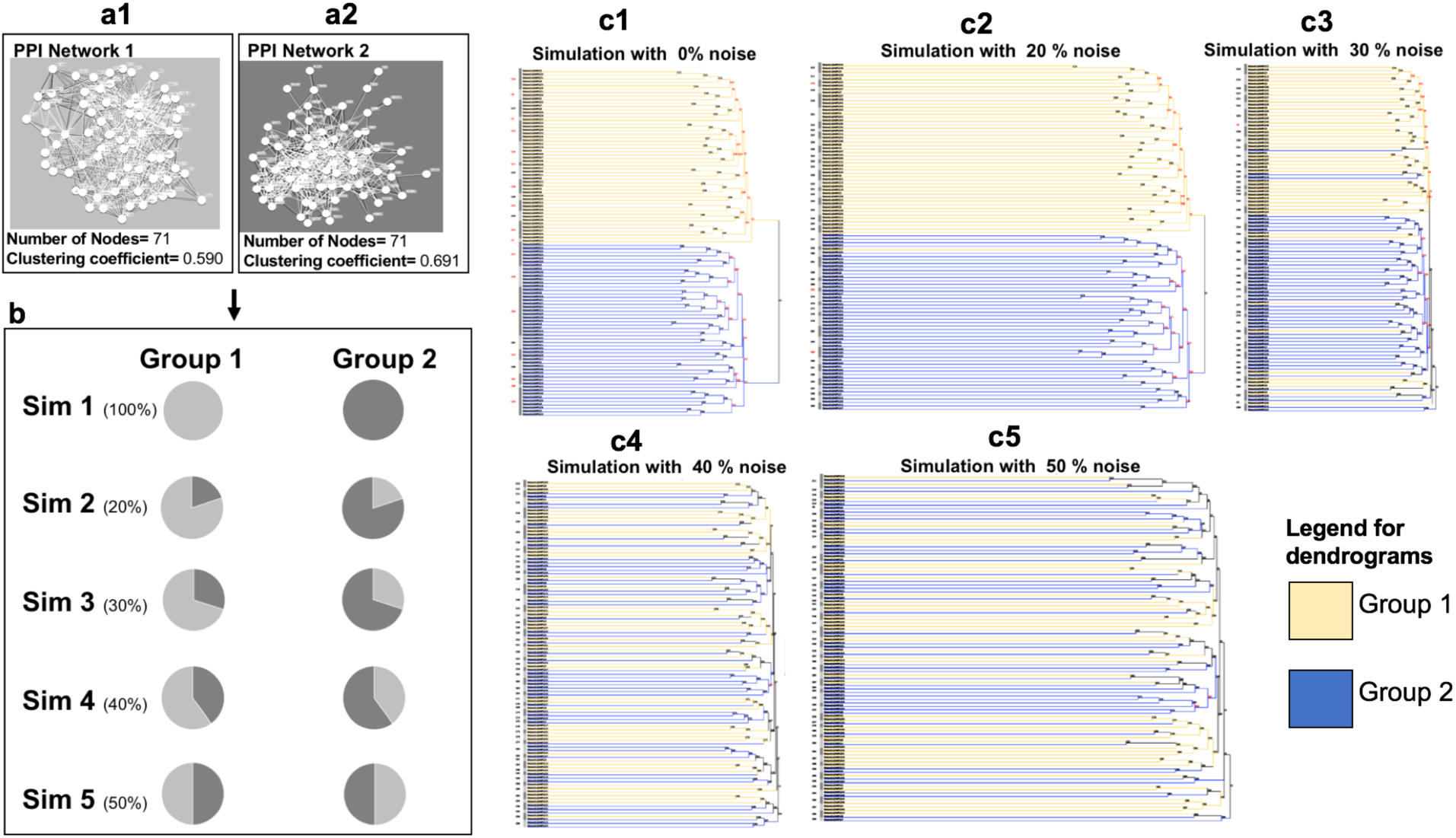
Simulations with Proteinarium on noisy data. **(a1 and 1a2)** STRING database was used to obtain PPI Networks 1 and 2, respectively. Both networks contain 71 proteins and their clustering coefficients were 0.590 and 0.691. There are no shared proteins between these two networks. **(b)** Simulated data was generated for Group1 and Group2, each with 50 samples. For each sample in Group 1, 10 genes were randomly selected from PPI. To add noise to the simulated data, for each sample in Group 1 a percentage of their genes were randomly replaced with genes from PPI Network 2 (20%, 30%, 40% and 50%). Similarly, for samples in Group 2, a percentage of genes were replaced at random with genes from PPI Network 1. This is represented by a series of pie chart schematics, showing the different amount of “noise” added to each sample in the various simulations (S1-S5). **(c1-c5)** Representative dendrograms reflecting the output of Proteinarium for the simulations with 0%, 20%, 30%, 40% and 50% noise respectively. Group 1 samples are represented in yellow, and Group 2 samples were represented in blue. The five dendrograms show the clustering of the samples based on their network similarities.

### Running Proteinarium on a Use Case

We implemented Proteinarium on a previously published genome wide expression study of preterm birth (26). The aim of the original study was to investigate maternal whole blood gene expression profiles associated with spontaneous preterm birth (SPTB, <37 weeks) in asymptomatic pregnant women. The study population was a matched subgroup of women who delivered at term (51 SPTBs, 114 term delivery controls). We used the gene expression microarray data generated from maternal blood collected between 27–33 weeks of gestation. The authors had performed univariate analyses to determine the differential gene expression associated with SPTB. We calculated z-scores for each gene in the preterm birth samples, using the mean of the control samples for reference (and vice-versa). For each sample, the genes were ranked according to their z-score and the top 50 genes were used as the sample’s gene list for input to Proteinarium. The dendrogram in Figure 3a demonstrates the clustering of the samples based on their network similarities.

**Figure 3.**
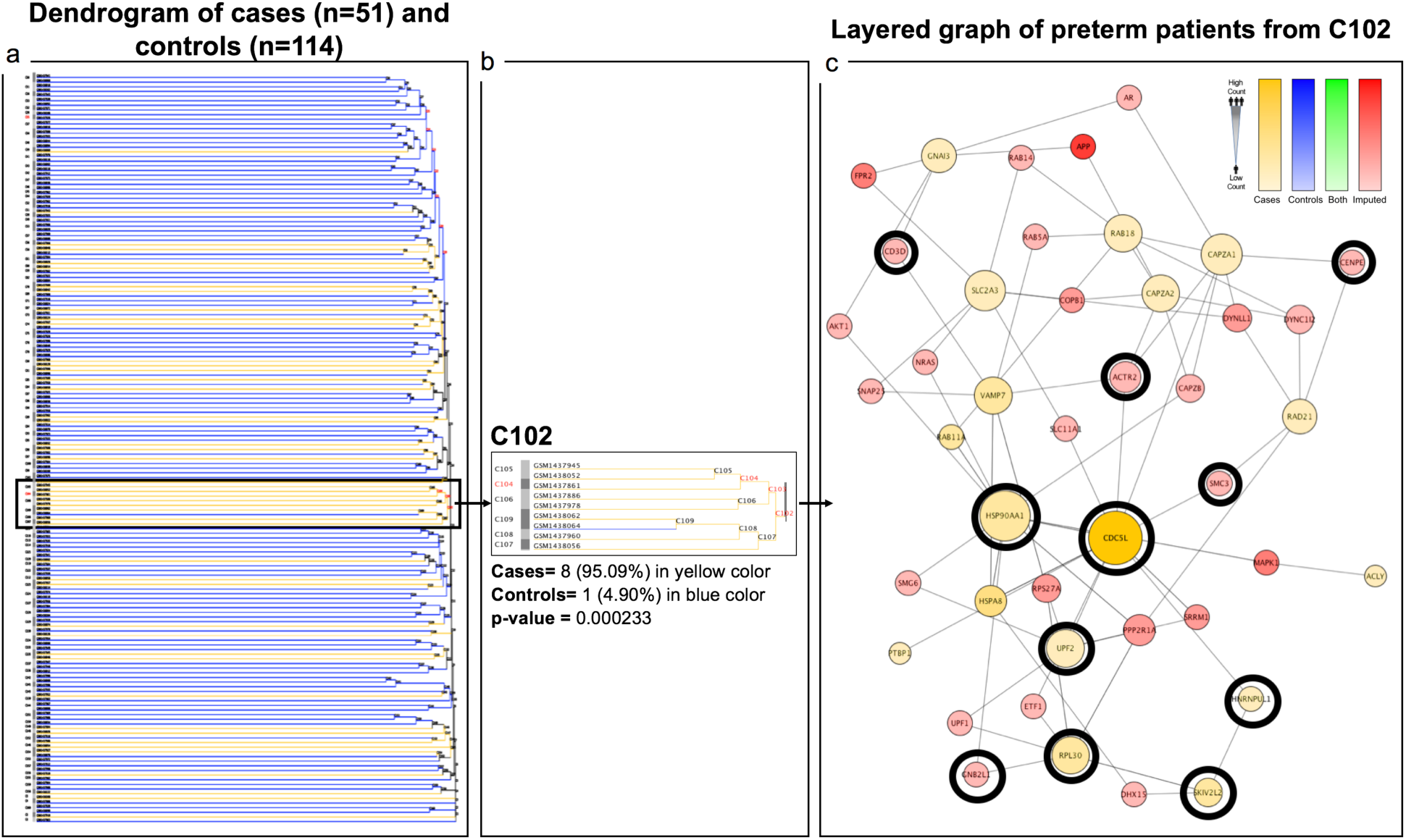
Proteinarium on a Use Case. Proteinarium was implemented on a previously published genome wide expression study of preterm birth. **(a)** Represents the dendrogram for cases and controls (51 SPTBs, 114 term delivery controls). **(b)** Cluster 102 (C102) represented in a zoom view. It contains significantly more preterm birth samples (n=8) than control samples (n=1). **(c)** Shows the layered PPI network of C102 for these preterm birth samples, consisting of 43 nodes. We found that 11 (in black circles) out of these 43 genes had previously been found to be nominally differentially expressed.

Cluster 102 contained significantly more preterm birth samples (n=8) than control samples (n=1). Figure 3c shows the layered PPI network for these preterm birth samples, consisting of 43 nodes. We found that 11 out of these 43 genes had previously been found to be nominally differentially expressed (26). Using over representation analysis, it was determined that this cluster is significantly enriched for preterm birth associated genes, both overall and amongst the inferred genes only (Chi-sq. 2 tailed p value=0.01). Additionally, the overall network density is 0.8, whereas the density for the subnetwork of the 11 PTB genes (Density = 0.16) is significantly greater(p=0.039). Using permutation testing, the probability of seeing a subnetwork of 11 nodes with a density of .16 or greater is less than 5%. Thus, the output from Proteinarium confirms and extends the results of this study. The results further support the validity of the assumptions underlying the design of Proteinarium.

## Discussion

Proteinarium is a multi-sample, protein-protein interaction tool built to identify clusters of samples with shared networks underlying complex disease. Using defined per sample seed genes, the Proteinarium pipeline takes input genes and converts them to protein symbols, which are then mapped onto the STRING PPI interactome. For each sample, its specific PPI network is built using Dijkstra algorithm by searching for the shortest path between each pair of protein inputs. The similarities between all subjects’ PPI Networks is calculated and used as the distance metric for clustering samples. In complex genetic diseases, Proteinarium can identify subsets of samples for which their shared networks differ between cases and controls.

In order to test Proteinarium, we simulated several data sets, each with a varying percentage of noise added to the data. With no noise, we confirmed full power to distinguish cluster 1 samples from cluster 2 samples (with 50 samples and input gene lists of size 10). As more noise was added to the data sets, the ability to discriminate between groups became more difficult, with the power decreasing from 100% down to 15% for 40% noise. We also validated that for the null hypothesis (i.e. there is no difference between Group 1 and Group 2) there was no power to discriminate with high confidence. This simulation was useful, first as a proof of concept that the algorithm performs as it should for both the most extreme as well as the null case. It also serves as a reference for users who might have prior information regarding the complexity or heterogeneity of their phenotypes.

We also implemented Proteinarium on a previously published genome wide expression study of a complex disease. As already mentioned, the aim of that study was to investigate maternal whole blood gene expression profiles associated with spontaneous preterm birth (SPTB, <37 weeks) in asymptomatic pregnant women. We ran Proteinarium on this data set, using the genes that had the highest Z-scores comparing SPTB cases with term controls as input. We found one Cluster for which the “CASE” cohort was highly represented. This cluster included 8 out of the 47 SPTB cases. We analyzed the layered PPI networks of these 8 subjects and found that there was significant over-representation of genes that had been previously found by the authors to be nominally significantly differentially expressed (26). Additionally, the network of these genes was denser than that of the whole network and had a high density that was unlikely to occur by chance alone. These findings support the validity of the assumptions underlying the design of Proteinarium and lend validity to the concept that the genetic architecture of complex disease is characterized by subgroups of patients that *share* variants in genes in specific networks and pathways which are sufficient to give rise to a phenotype.

One of strengths of Proteinarium is the ability of the user to configure all parameters throughout the pipeline. For example, one of the configuration options is the maximum path length (MPL). MPL is the maximum number of vertices (or imputed genes) within any given path. Thus, a user defined MPL of 2 would allow single genes to be imputed between any two of the seed genes. This parameter allows for flexibility in building a network with varying degrees of distant neighbors. With a MPL of 2 or greater, proteins that connect a pair of the input nodes will be added as imputed nodes to the network. This allows for inference(s) on genes which may have not originally been considered as relevant based on the results used to generate the input data. Once the individual networks are generated and pairwise similarities are calculated, hierarchical clustering is used to cluster the samples. The output of this analysis is displayed as a dendrogram. In addition to the image of the dendrogram (.png), Proteinarium provides the dendrogram structure in Newick tree format. This allows users to visualize the dendrogram on other platforms like Dendroscope or ITOL (23, 24) and for subsequent analysis with other software. An additional output is the Cluster Analysis File, which provides detailed information about clustering results in a tabular format (.csv). This file contains the clustering coefficient of the layered graph for that cluster, the number of samples in the cluster, the p-values for statistical abundance of group type, and the average distance (height) for the branches of the dendrogram for each cluster. This cluster is also available on the fly on the command line screen when a cluster is selected.

In addition to its use as an analytic tool, Proteinarium provides useful visualizations. The dendrogram, allows the user a global view of the distribution of samples into clusters and the group coloring allows for identification of patterns related to phenotypic class. Additionally, as mentioned above, users can select any branch of the dendrogram and display the layered PPI network of the samples within that cluster. This functionality enables users to visualize the layered networks of multiple samples. If a cluster contains samples from *both* phenotypic groups, then there are 5 possible networks created on the fly. These network images allow users to visually identify interesting aspects of the network, like the most connected genes, the most frequently appearing genes amongst the samples in the cluster, the distribution of genes originated from each phenotypic group as well as those that are imputed. Proteinarium also provides an output file for these networks. The file includes information about the genes in the network, if they are imputed or not and the number of samples that contain each gene. Additionally, Proteinarium provides a PPI interaction file for each network. This allows users to analyze the network(s) of interest on other platforms such as Cytoscape or Gephi (27, 28).

### Future applications

The current release of Proteinarium uses protein-protein interaction data from a single repository, the STRING database. We plan to extend the resource options in the next release by allowing the PPI information data to be derived from any IMEX consortium that the users choose (29). In addition, we plan to allow PPI from other organisms in Proteinarium, e.g. mouse. In parallel, these extensions will grow by implementing a user interface. Users will be able to visualize the networks by clicking the on the dendrogram rather than using the command line options.

## Conclusion

In conclusion, we have created a multi-sample protein-protein interaction network tool to support analysis and visualization of single or paired samples. The tool allows investigators to address important questions from their high throughput data for a variety of disease phenotypes based on their associated PPI networks. In addition, Proteinarium provides several different, user-defined outputs with more than 30 configurable options.

## Availability and requirements

**Project name:** Proteinarium

**Project home page:** https://github.com/Armanious/Proteinarium

**Operating system(s):** Linux, Mac OS X, Windows

**Programming language:** Java

**Other requirements:** none

**License:** GNU Affero GPL (Version 3)

**Any restrictions to use by non-academics:** None

## List of abbreviations

C: Cluster
HGNC: HUGO gene nomenclature committee
PPI: Protein-protein interactions
ITOL: **Interactive tree of life**
UPGMA: Unweighted pair group method with arithmetic mean
LG: Layered graph

## Declarations

### Ethics approval and consent to participate

Not applicable.

### Consent for publication

Not applicable.

### Availability of data and material

The sample datasets used during the development of current software and the latest version can be freely downloaded at https://github.com/Armanious/Proteinarium

### Competing interests

The authors declare that they have no competing interests.

### Funding

This work was supported by the National Institutes of Health (grants 5P20GM109035-04, 5P30GM114750) and the Kilguss Research Core at Women & Infants Hospital.

### Authors’ contributions

D.A., A.U., J.S. developed the algorithm. D.A. wrote the software. S.I., and J.P. provided guidance and contributed to the features of the software. D.A., J.S. and A.U. wrote the manuscript with critical feedback and input from all authors. A.A did the analysis on the simulated data sets. G.F.T. worked on the use case dataset. A.T.D contributed into statistics and analysis approaches of the study. J.P., J.S. and A.U. edited the manuscript. A.U. conceptualized the project and supervised the study.

## Acknowledgements

We thank Stephen Kidd from the Department of Classics at Brown University for naming the tool. We thank the Center for Computation and Visualization (CCV) and Computational Biology Core at Brown University.

## Notes

#### Summary of Updates

In this updated version, we included the extended analysis of simulations and we added use case for Proteinarium analysis. The overall text was modified accordingly as well.

https://github.com/Armanious/Proteinarium

