## Supplemental File for "Proteinarium: Multi-Sample Protein-Protein Interaction Analysis and Visualization Tool"

**Supplementary Material**

### 1. Visualizing “Local” Clusters

#### 1a. Constructing a layered graphs vi and v:

Given two layered graphs $LG_1$ and $LG_2$, we define the subtraction operation $LG_1 - LG_2$ to be: for each vertex $v \in LG_1$, if either (1) $v \notin LG_2$ or (2) the scaled count of $v$ in $LG_1$ minus the scaled count in $LG_2$ is greater than 0.

```
LayeredGraph subtract(LG1, LG2):
G = {vertices, edges}
for v in LG1.vertices:
    If LG1.count(v) > LG2.count(v):
        G.vertices.add(v, LG1.count(v) - LG2.count(v))
for e in LG1.edges:
    If e.source in vertices and e.target in vertices:
        G.edges.add(e)
remove extraneous vertices from G.vertices according to G.edges
return G
```

For the purposes of visualization of the newly constructed graph $G$, we scale the count annotations of $LG_1$ and $LG_2$ by the parameter $\rho$ as defined above.

#### 1b. Visualizing and annotating the layered graph: Force Directed Layout

For each vertex $v_i \in LG$, we assign its radius $r_i$ linearly according to degree between some specified minimum and maximum radii. Let $d_{ij}$ be the Euclidean distance between the center points of two vertices, $v_i, v_j$. For any neighboring pair of vertices $v_i, v_j$, we define the following spring-based attractive force: $F_A = \lambda_A d_{ij}$. For every pair of vertices $v_i, v_j$, we define the following electrostatic repulsive force: $F_R = -\frac{\lambda_R r_i r_j}{d_{ij}^2}$. We then apply the force $F_A + F_R$ to $v_i$ at the angle between $v_i$ and $v_j$. Doing this for all vertices constitutes 1 iteration, and we continue iterations until either [maxIterations] or [maxTime] is exceeded, or when the total distance updates over all vertices is less than [deltaThreshold].
